# Plasticity of mitochondrial function safeguards phosphorylating respiration during *in vitro* simulation of rest-phase hypothermia

**DOI:** 10.1101/2022.05.25.493417

**Authors:** Carmen C. García-Díaz, Imen Chamkha, Eskil Elmér, Andreas Nord

## Abstract

Many animals downregulate body temperature to save energy when resting (rest-phase hypothermia). Small birds that winter at high latitude have comparatively limited capacity for hypothermia and so pay large energy costs for thermoregulation during cold nights. Available evidence suggests this process is fuelled by adenosine triphosphate (ATP)-dependent mechanisms. Most ATP is produced by oxidative phosphorylation in the mitochondria, but mitochondrial respiration can be lower during hypothermia because of the temperature-dependence of biological processes. This can create conflict between increased organismal ATP demand and a lower mitochondrial capacity to provide it. We studied this in blood cell mitochondria of wild great tits (*Parus major*) by simulating rest-phase hypothermia *via* a 6°C reduction in assay temperature *in vitro*. The birds had spent the night preceding the experiment in thermoneutrality or in temperatures representing mild or very cold winter nights. Night temperature did not affect mitochondrial respiration. Across treatments, endogenous respiration was 14% lower in hypothermia. This did not reflect general thermal suppression because phosphorylating respiration was unaffected by thermal state. Instead, hypothermia was associated with a threefold reduction of leak respiration, from 17% in normothermia to 4% in hypothermia. Thus, coupling of total respiration to ATP production was 96% in hypothermia, compared to 83% in normothermia. Our study shows that thermal insensitivity of phosphorylation combined with short-term plasticity of leak respiration may safeguard ATP production when endogenous respiration is suppressed. This casts new light on the process by which small birds endure harsh winter cold and warrants future tests across tissues *in vivo*.

## INTRODUCTION

Endothermic animals, such as mammals and birds, use considerable amounts of energy to regulate a high and relatively stable body temperature across a wide range of air temperatures. The associated energy cost of thermoregulation can sometimes become prohibitively high, for example when food is scarce or when weather conditions deteriorate. Many small to medium-sized endotherms use rest-phase hypothermia, a regulated reduction of resting body temperature below normal levels, to mitigate such energetic challenges. This may reduce metabolic demands by 50% or more [1]. It follows that most animals using rest-phase hypothermia have high metabolic intensity, lose heat quickly and rely on high-energy or ephemeral food sources [2]. This fits the description of “the little bird in winter” [3], i.e., small (< 20 g) songbirds that are year-round residents at high latitudes where they routinely meet environmental temperatures 50-60 °C below body temperature during the cold season. Yet, most birds in northern forests, such as tits and chickadees, typically show daily variation in body temperature within 5 °C [e.g. 4, 5, 6] and generally do not exceed 7-8 °C even in severe cold [7-9, but see 10 for an exception]. Theoretical models predict that even such moderate body temperature reduction can carry survival benefits [11, 12]. However, the modest extent of hypothermia means that the little bird in winter can never forego a significant increase in energetically costly heat production to counter winter cold, even when maximally hypothermic [e.g. 13].

Most of the energy used by the working body is in the form of adenosine triphosphate (ATP) that is produced mainly by oxidative phosphorylation in the mitochondria [14]. Since more energy is needed to stay warm in winter, it is not surprising that many endotherms upregulate mitochondrial content and/or respiration rate in key metabolic tissues when cold exposed, such as in the wintertime [15-18, e.g. 19]. However, little is known about mitochondrial adjustment in response to shorter-term variation in air temperature, for example during a sudden drop in night-time temperature coincident with clearing of an overcast sky. Moreover, a potential problem for a hypothermic animal is that the activity of the respiratory chain could be slower in a colder body [e.g. 20], causing a conflict between increased organismal demand for fuel but reduced mitochondrial capacity to provide it. However, not all the proton motive force is used to produce ATP since some protons backflow into the mitochondrial matrix bypassing ATP synthase. This is mediated by three principal mechanisms. Physiological properties maintain a basal level of proton conductance of the mitochondrial membrane that can be further up-or downregulated under the influence of uncoupling proteins [21]. A functionally similar phenomenon occurs when protons spontaneously “slip” back through the proton pumps of the electron transport chain, though the contribution of slippage to total uncoupling is probably low [reviewed by 21]. Compensation for these processes make up what is known as leak respiration. This constitutes some 20-25% of resting metabolic rate in endotherms [22, 23], meaning there is scope to adjust the coupling of electron transport to ATP production according to the animal’s need. For example, increased mitochondrial uncoupling in brown adipose tissue (BAT) is the basis for non-shivering heat production in mammals [14]. In contrast, birds lack BAT and with few exceptions are thought to produce heat using ATP-dependent shivering in skeletal muscles [24, 25; but see 26-27 for exceptions]. Thus, in birds it can be speculated that acute cold exposure could be associated with improved coupling, to make ATP production more effective for a given respiration rate, which has been suggested for other energy demanding processes such as fasting [e.g. 28-31]. This could secure adequate fuel delivery even when cell respiration is reduced, such as when heat production must remain high, but rest-phase hypothermia causes the mitochondria to work at slower pace. There is support for this compensation hypothesis from studies on ectotherms [32-34; but see 35, 36 for exceptions]. Similar studies on endothermic mammals suggest that while phosphorylating respiration is often suppressed in hypometabolic animals, non-phosphorylating (i.e., leak) respiration is sometimes reduced, sometimes unaltered, and sometimes increased [reviewed by 37].

Previous endotherm studies have largely been performed in mammals during hibernation or in deep torpor; thermal states characterised by substantial metabolic downregulation [1] beyond what is feasible in most birds [but see 38]. Thus, previous data are not easily extrapolated to the problems faced by the little bird in winter that must increase energy expenditure sharply to counter cold temperature even when hypothermic and must do so using ATP-dependent thermogenesis. Hence, we studied mitochondrial responses to both cold environments and rest-phase hypothermia in the great tit (*Parus major* L.) – a small (16-20 g) songbird that winters at high latitude and show pronounced diel variation in body temperature [4, 39]. We exploited the fact that birds contain functional mitochondria in their red blood cells. Previous studies have found that avian blood cell mitochondria respond in the short-term to variation in energetic demands [40-42], over longer time periods in line with age-related metabolic senescence [43] and according to environmental context [18]. There is also evidence suggesting blood cell respiration is correlated to that in in key tissues for heat production (skeletal muscle) and energy management (heart, kidney, brain) in mammals and birds [44-46], all of which show pronounced seasonal acclimatisation to winter conditions in small, northern birds [47]. This emphasises the need for further studies exploring the utility of blood cell respirometry as a minimally invasive method for assessing mitochondrial responses to environmental stressors. On this basis, we, firstly, tested if the immediate thermal environment of the bird impacted mitochondrial function by manipulating the night preceding measurements to be representative of mild or very cold winter temperature, or thermoneutrality. Then, we performed an *in vitro* simulation of rest-phase hypothermia, to test if lower body temperature caused a reduction in mitochondrial respiration and if this was associated with any changes in the efficiency of ATP-production. We expected that low night-time temperature would be associated with increased mitochondrial respiration rate to facilitate ATP-dependent heat production. In contrast, we predicted that lower assay temperature would cause thermal suppression of respiration traits. However, if better coupled mitochondria serve as an adaptation to safeguard ATP production when cold ambient temperature increases organismal-level demand for ATP, or when lower body temperature reduces overall energy expenditure of the cell, we expected tighter coupling of electron transport to ATP production both in response to lower air temperature at night and during *in vitro* hypothermia.

## MATERIALS AND METHODS

The experimental design and timeline are outlined in Fig. 1.

**Figure 1.**
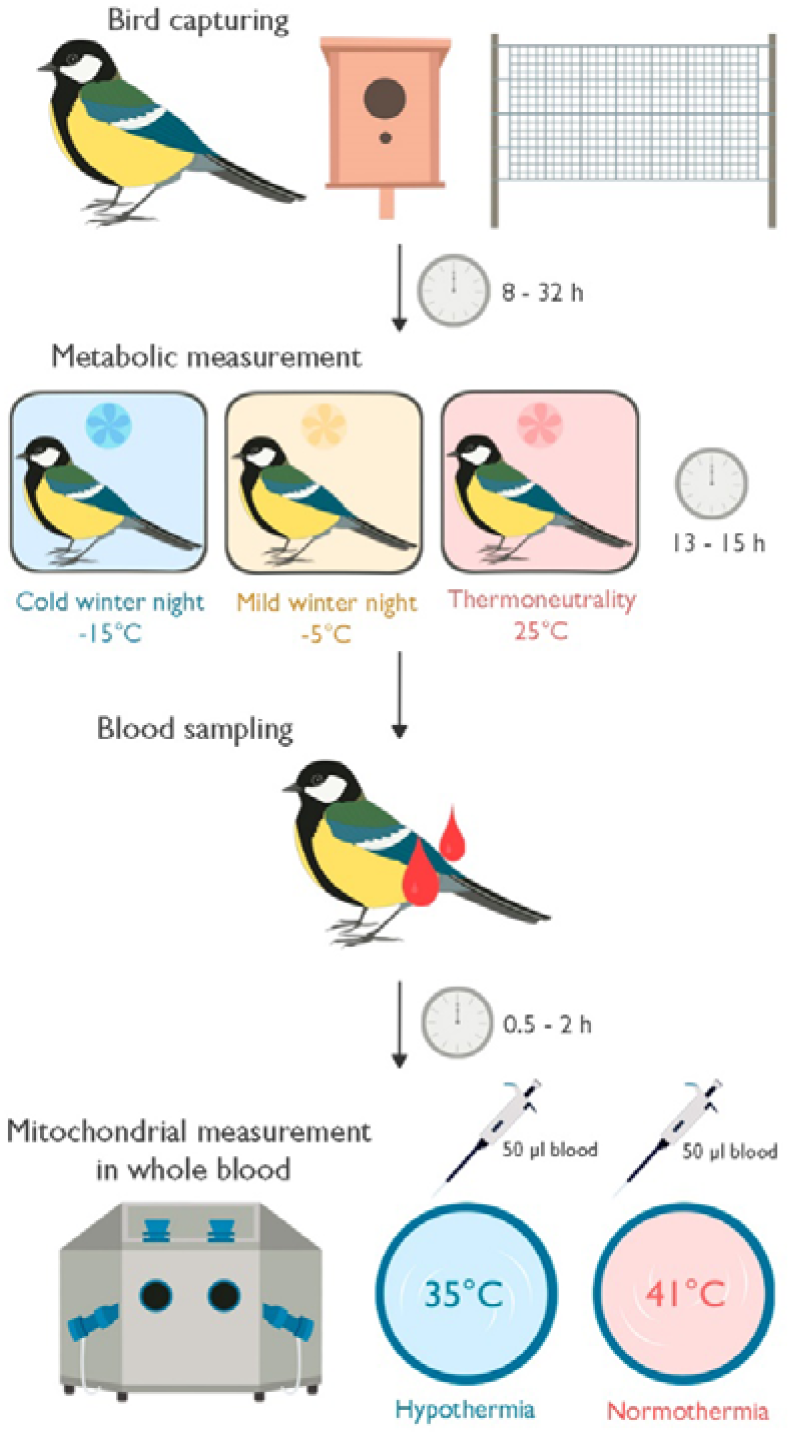
Overview of the experimental design and timeline of events when studying the effect of thermal state (normothermia or hypothermia) on functional properties of mitochondria in avian blood cells. Studies were performed using wild great tits (*Parus major*) that wintered outside the city of Lund in southernmost Sweden.

### Study site and animals

Great tits were caught at two sites in the vicinity of Lund in southernmost Sweden – Räften (n = 37) and Vomb (n = 21) – between January and February 2021. Räften (55° 43’N, 13° 17’E”) is an oak (*Quercus robur*) dominated 32 ha woodlot surrounded by arable fields. The Vomb site (55° 39’N, 13° 33’E) is part of a pine (*Pinus sylvestris*) plantation with a rich deciduous understory composed mainly of sallow (*Salix* spp.), birch (*Betula pendula*) and rowan (*Sorbus aucuparia*). Birds in Räften were caught in mist nets close to feeders baited with sunflower seeds that were erected 1.5 m above ground in January 2021. The birds in Vomb were caught shortly before sunrise when roosting in nest boxes. Four to eight birds were caught each day of the study and were brought to Lund University within 2 h of capture, where they were alternately allocated to temperature treatment groups in batches of four. We housed the birds in cages measuring 77 × 44 × 77 cm (length × width × height) at 5 °C under simulated natural photoperiod and were provided sunflower seeds, peanuts, and water *ad libitum*. One-two days after capture the birds were implanted with a temperature-sensitive PIT tag (BioTherm13, Biomark, Boise, ID, USA) into the intraperitoneal cavity under local anaesthesia (5% lidocaine) as part of a different study. Eight to 34 h later, the birds were put in a 1 L hermetically sealed glass container ventilated with dry air at 400-500 ml / min (recorded using a FB-8 mass flow meter, Sable Systems, Las Vegas, NV, USA) for measurement of resting metabolic rate (RMR; as oxygen consumption recorded in carbon dioxide-free air using a FC-10 oxygen analyser, Sable Systems) and body temperature (recorded using a custom-built multiplexed antennae system from BioMark) in a climate chamber (Weiss Umwelttechnik C180 [Reiskirchen, Germany]) at thermoneutrality (25 °C; n = 20) [13], during a simulated mild winter night (5 °C: n = 19) and during a simulated very cold winter night (−15 °C; n = 19) for 13-15 h. Four birds were measured each night. Mean RMR was 0.36 ± 0.01 W (mean ± s.e.m.), 0.51 ± 0.01 W and 0.68 ± 0.02 W in +25, +5 and -15 °C, respectively. Mean nightly body temperature was 37.9 ± 0.3 °C (range of means: 35.2 to 40.1 °C; minimum: 33.8 °C), 39.3 ± 0.1°C (range of means: 37.6 to 40.2 °C; minimum: 37.0 °C), and 39.2 ± 0.1 °C (range of means: 38.0 to 40.3 °C; minimum: 37.6 °C), in the cold, mild, and thermoneutral night conditions.

Air temperature in Lund (10-40 km from the study sites) ranged -14 °C to 7 °C during the study period and photoperiod ranged 8:20 h to 10:20 h.

### Blood sampling and mitochondrial measurement in whole blood

In the morning after the metabolic measurements, a 100-150 µl blood sample ≤10% of estimated total blood volume was obtained from the jugular vein using a heparinized syringe within 5 min of removing the bird from the metabolic chamber. The sample was stored at 10-12 °C in 2 mL EDTA (ethylenediaminetetraacetic acid) tubes (BD Vacutainer^®^, Franklin Lakes, NJ, USA) without agitation until analysed 0.5-2.5 h later. All birds were released at the site of capture within 2 h of blood sampling.

We measured mitochondrial respiration in whole blood using an Oxygraph O2k high resolution respirometer (Oroboros Instruments, Innsbruck, Austria), following the protocol described in Nord, Chamkha & Elmér [48]. To test how rest-phase hypothermia affected mitochondrial function, we ran duplicate samples from the same individual simultaneously in each of two thermal states: a representative normothermic daytime bird body temperature (41 °C; henceforth “normothermia”) and a hypothermic night-time body temperature in the lower range of the measured natural variation (35 °C; henceforth “hypothermia”) [49] using two respirometers. These were calibrated at assay temperature (i.e., 35 °C or 41 °C) daily, and we changed instrument/assay temperature combinations between days to avoid confounding experimental results by any variation pertaining to the specific instruments. The main purpose of our study was to demonstrate the presence of a putative mitochondrial response to acute body temperature change. To achieve this goal, we purposedly selected a low but biologically relevant hypothermic assay temperature, under the assumption that any phenotypic response would be linear over the range of *in vitro* temperatures considered.

The blood samples were allowed 5-10 min on a rotating mixer at room temperature before the start of the assay. Then, we added 50 µL sample to 1.95 mL MiR05 respiration medium (0.5 mM EGTA, 3 mM MgCl_2_, 60 mM K-lactobionate, 20 mM taurine, 10 mM KH_2_PO_4_, 20 mM Hepes, 110 mM sucrose, free fatty acid bovine serum albumin (1 g L^-1^), pH 7.1) that was contained within the respirometer chambers and so was pre-equilibrated at assay temperature. The final volume contained in the respirometer chamber was, thus, 2 mL.

Mitochondrial respiration rate was measured as the rate of decline in O_2_ concentration in the chamber. Once the O_2_ consumption had stabilized upon closing the chamber, baseline O_2_ consumption on endogenous substrates (‘ROUTINE’) was recorded for 2-3 min. Then, we assessed mitochondrial function by sequential addition of three mitochondrial agonists and antagonists. Firstly, we added the ATP-synthase inhibitor oligomycin at a final concentration of 5 µmol L^-1^, which blocks the process of oxidative phosphorylation (‘OXPHOS’). Thus, the remaining O_2_ after oligomycin addition is attributed to proton leak (‘LEAK’). Next, we added the mitochondrial uncoupler FCCP (carbonyl cyanide-p-trifluoro-methoxyphenyl-hydrazone), which abolishes the proton gradient across the inner mitochondrial membrane, causing the electron transport system to work at maximum capacity to restore the gradient (‘ETS’). Because an excessive concentration of FCCP inhibits mitochondrial respiration and can lead to acidification of the cell cytoplasm [14], we first added 1.5 µL of 1 mmol L^-1^ FCCP and then titrated FCCP in 0.5 µL increments until maximum O_2_ consumption was reached. The final concentration ranged 1-1.75 µmol L^-1^. Finally, we added 5 µmol L^-1^ of antimycin A, which blocks Complex III of the electron transport chain. Therefore, the remaining O_2_ consumption after antimycin A addition is due to non-mitochondrial oxygen utilisation.

### Data handling and statistical analyses

We excluded five of the 57 duplicate samples measured at 41 °C since they were not inhibited by the addition of oligomycin. Thus, the final data set consisted of 57 observations at 35 °C and 52 observations at 41 °C. We then calculated mitochondrial respiration states and flux control ratios (FCR; i.e., ratios between the respiratory states) following Gnaiger (2020) [50]. We considered three FCRs: 1) E-R control efficiency (1 - ROUTINE / ETS), which measures how much excess capacity there is in the electron transport system until maximum respiration is reached); 2) R-L control efficiency ((ROUTINE - LEAK) / ROUTINE), which measures the proportion of endogenous respiration that is channeled towards oxidative phosphorylation; 3) E-L coupling effiency (1 - LEAK / ETS) which is indicative of mitochondrial efficiency in the context of coupling between electron transport and ATP production in a stimulated cellular state. Derivation and definition of all respiration traits and FCRs are detailed in Table 1.

**Table 1.** Calculation method and definition of mitochondrial respiration states and flux control ratios. All traits were derived based on changes in O_2_ consumption rate in response to the addition of mitochondrial agonists and antagonists during the assay as described in the main text.

| Parameter | Calculation method | Definition |
| --- | --- | --- |
| ROUTINE | $R_{\text{Baseline}} - R_{\text{Antimycin A}}$ | Baseline O<sub>2</sub> consumption during oxidative phosphorylation under endogenous cellular conditions. |
| OXPHOS | $R_{\text{Baseline}} - R_{\text{Oligomycin}}$ | O<sub>2</sub> consumption used to drive oxidative phosphorylation where ATP is produced. |
| ETS | $R_{\text{FCCP}} - R_{\text{Antimycin A}}$ | O<sub>2</sub> consumption when the electron transport system is operating at its maximum capacity. |
| LEAK | $R_{\text{Oligomycin}} - R_{\text{Antimycin A}}$ | O<sub>2</sub> consumption needed to compensate for proton leakiness of the mitochondria. |
| E-R control efficiency | $1 - (\text{ROUTINE} / \text{ETS})$ | Proportion of ETS maximum working capacity remaining under endogenous cellular conditions. |
| R-L control efficiency | $(\text{ROUTINE} - \text{LEAK}) / \text{ROUTINE}$ | Proportion of endogenous respiration used for ATP production via oxidative phosphorylation. |
| E-L coupling efficiency | $1 - (\text{LEAK} / \text{ETS})$ | Tightness of electron transport during ATP production in a stimulated cellular state. |

All statistical analyses were performed using R version 4.1.2. To test how a sudden change in night temperature affected mitochondrial respiration the morning after, we used linear models (lm in R base) to test how each respiration state and FCR assayed at normothermic temperature (i.e., 41°C) was affected by preceding night temperature. We used linear mixed models (lmer in the lme4 package) [51] to test how respiration traits and FCRs were affected by thermal state (i.e., hypo-or normothermia) using bird ID as a random factor to account for repeated measurements. To test if the phenotypic response to a change in thermal state differed depending on the depth of hypothermia during the preceding night, we included mean nightly body temperature and body temperature × thermal state in these models. Significances were assessed using *F*-tests with degrees of freedom based on the Kenward-Roger approximation, implemented using the KRmodcomp function in the pbkrtest package [52]. The significance of the random term (i.e., bird ID) in final models was assessed using log likelihood tests using the drop1 function in base R. The interaction was removed when non-significant (*P* > 0.05), but all main effects were retained. Predicted values and standard errors (s.e.m.) for the lmer models were calculated using the emmeans function in the emmeans package [53]. For the tests of night temperature effects, estimates and their s.e.m. were taken from the linear model output.

## RESULTS

A representative normothermic experiment is presented in Fig. S1. All details of the statistical tests pertaining to effects of night temperature are presented in Table S1, and those pertaining to the effects of thermal state and body temperature are presented in Table S2.

### Effects of preceding night temperature

All but one of the mitochondrial respiration metrics were unaffected by temperature of the preceding night (all *P* > 0.16; Table S1). There was a tendency for R-L control efficiency, which measures how much of the ROUTINE respiration that is channelled towards ATP production, to vary with night temperature (*P* = 0.063). Specifically, 86% of endogenous respiration was used to drive oxidative phosphorylation after a night in -15°C, which was higher than after a night at +5°C (81%) but similar to a night in +25°C (83%) (Table S1).

### Effects of thermal state and nightly body temperature

There were profound changes when assay temperature changed from a normothermic to a hypothermic thermal state. ROUTINE respiration was 14 % higher in normothermia (0.567±0.019 pmol O_2_ s^-1^ µl^-1^) than in hypothermia (0.499±0.019 pmol O_2_ s^-1^ µl^-1^) (*P* < 0.0001) (Fig. 2A), i.e., endogenous respiration was lower during *in vitro* hypothermia. However, neither OXPHOS nor ETS were affected by thermal state (*P* = 0.7 and 0.6, respectively) (Figs. 2B-C). Instead, the drop in ROUTINE was mediated by a pronounced reduction in LEAK respiration (*P* < 0.0001), which was more than three times higher during normothermia (0.096±0.004 pmol O_2_ s^-1^ µl^-1^) compared to hypothermia (0.023±0.004 pmol O_2_ s^-1^ µl^-1^) (Fig. 2D).

**Figure 2.**
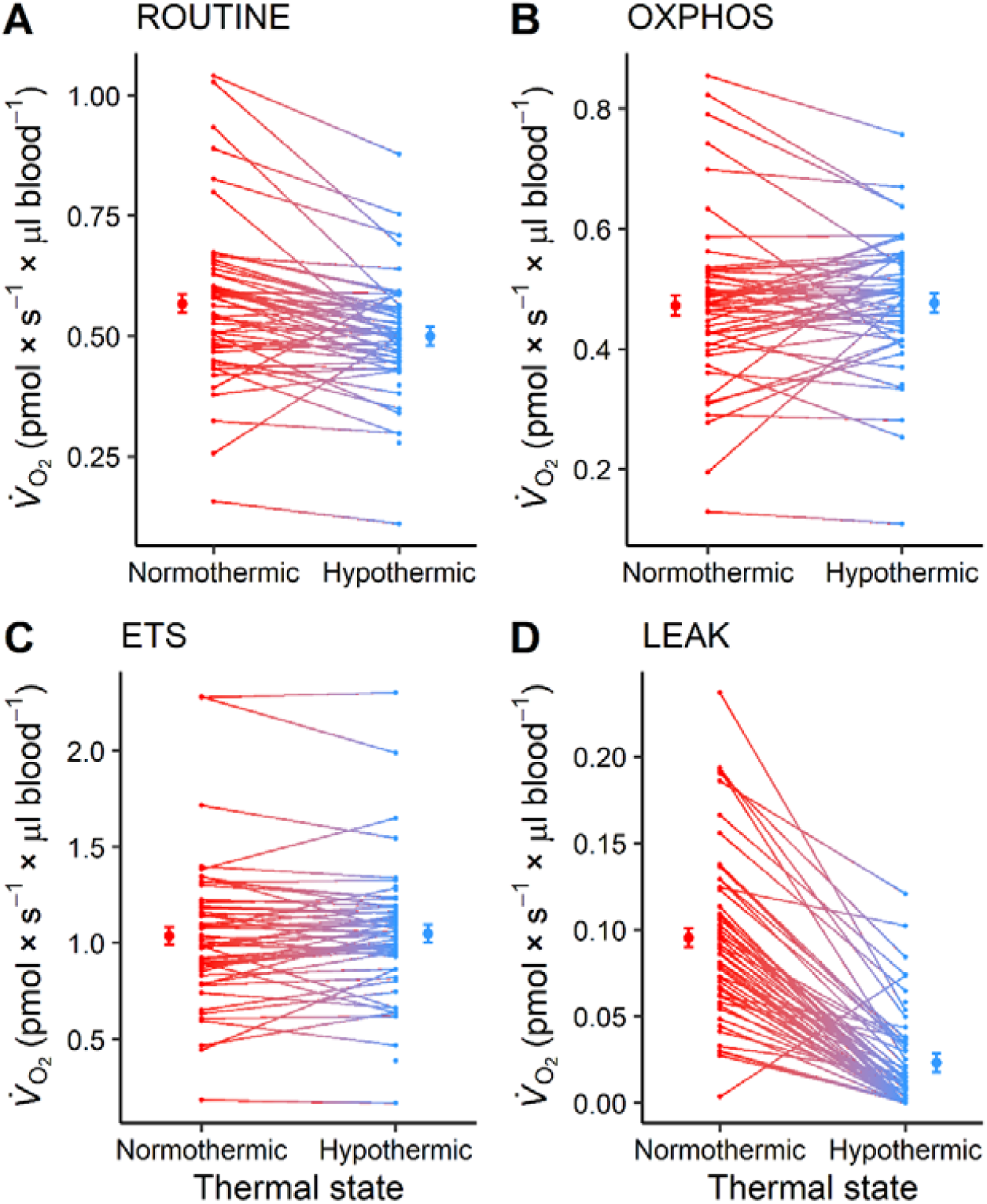
Mitochondrial respiration traits in whole blood of winter-adapted, wild, great tits (*Parus major*) at a representative daytime normothermic body temperature (41 °C) and a representative night-time hypothermic body temperature (35 °C). The same individuals were measured in both thermal states. Solid plotting symbols with error bars represent model-predicted means ± s.e.m. Gradient lines show individual responses. Sample sizes and statistics are reported in Table S2. (A) baseline (“ROUTINE”) oxygen consumption during oxidative phosphorylation. (B) oxygen consumption devoted to ATP production alone (“OXPHOS”) (C) maximum working capacity of the electron transport system (“ETS”). (D) the part of ROUTINE used to offset proton leak across the inner mitochondrial membrane (“LEAK”).

E-R control efficiency was lower during hypothermia (*P* < 0.0001), meaning a hypothermic bird had more mitochondrial reserve capacity compared to a normothermic bird (0.505±0.012 and 0.434±0.012, of 1, respectively) (Fig. 3A). The efficiency of oxidative phosphorylation was higher in the hypothermic state (i.e., R-L control efficiency increased) such that nearly all (96%) of endogenous respiration was used to produce ATP in hypothermia, compared to 83% in normothermia (Fig. 3B). In other words, LEAK accounted for nearly 17% of endogenous respiration during normothermia, but in hypothermia LEAK contributed only 4% to ROUTINE. E-L coupling efficiency (i.e., coupling of the ETS when maximal respiration was stimulated using FCCP) increased from 91% (0.907±0.004) in normothermia to 98% (0.978±0.004) in hypothermia (Fig. 3C). Thus, the electron transport chain was tighter in the hypothermic state.

**Figure 3.**
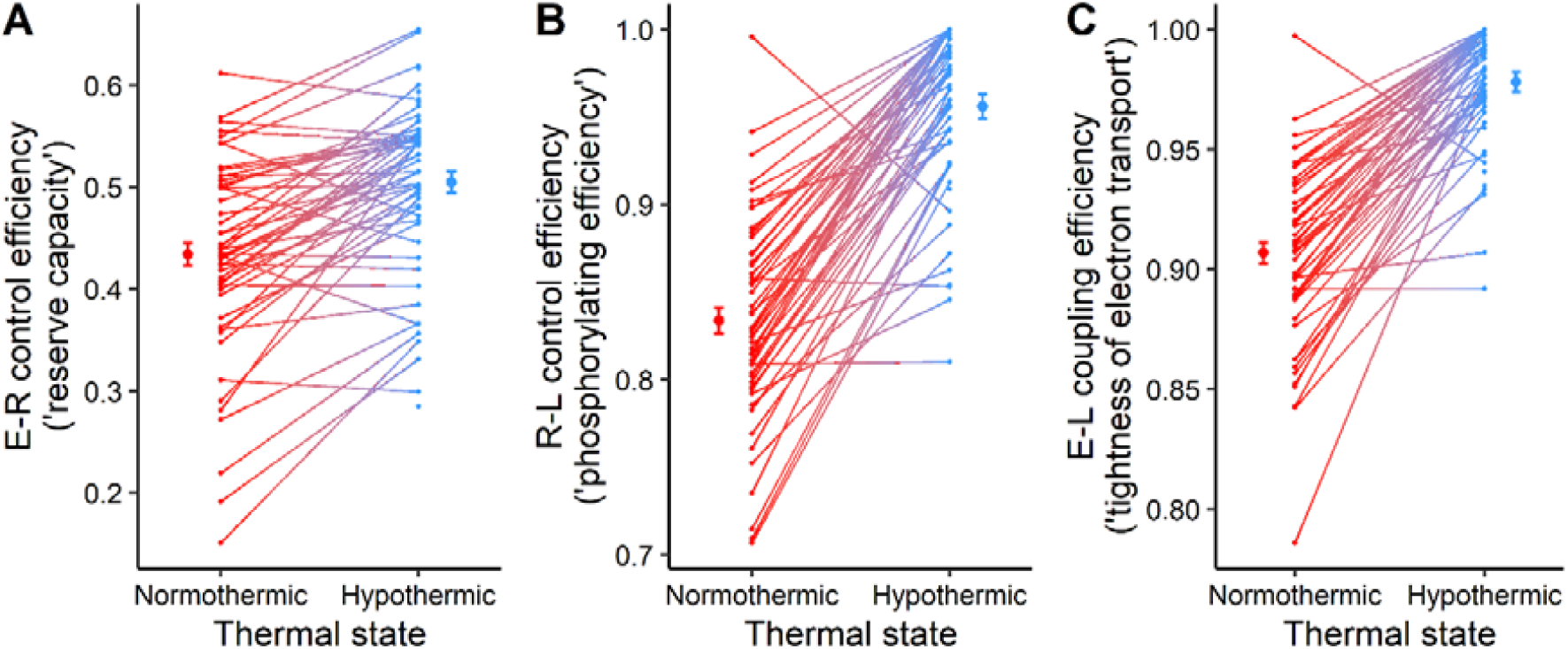
Control ratios for mitochondrial respiration in great tit whole blood in a normothermic (41 °C) and a hypothermic (35 °C) thermal state. (A) E-R control efficiency indicates how intensively the mitochondria were respiring under endogenous conditions (i.e., ROUTINE) relative to their maximum working capacity (i.e., ETS). For example, a value of 0.4 indicates that mitochondrial respiration can increase by 40% before reaching maximum. (B) The fraction of endogenous respiration that is used for oxidative phosphorylation where ATP is formed. (C) Coupling efficiency of electron transport when the ETS is stimulated to work at maximum capacity, where a value of 0 indicates a fully uncoupled system and a value of 1 a fully coupled system (i.e., where all respiration is towards phosphorylation). Samples were collected from wild birds and the same individual was measured in both thermal states. Solid plotting symbols with error bars represent model-predicted means ± s.e.m. Gradient lines show individual responses. Sample sizes and statistics are reported in Table S2.

Most mitochondrial respiration traits were unaffected by nightly mean body temperature, both as main effects and in interaction with thermal state (Table S2; Fig. 4). However, E-R control efficiency increased with increasing body temperature (by 0.041±0.010 per °C) across thermal states (Fig .4), suggesting that birds that were more hypothermic during the preceding night had mitochondria that respired at a greater proportion of maximum. There was also a thermal-state-dependent relationship between R-L control and body temperature, whereby oxidative efficiency decreased by 0.012±0.008 per °C in the normothermic state but was unaffected by body temperature in the hypothermic state. However, when tested in assay temperature-specific models, the linear regression between R-L control and mean night-time body temperature was not significant (hypothermia: *P* = 0.39; normothermia: *P* = 0.13).

**Figure 4.**
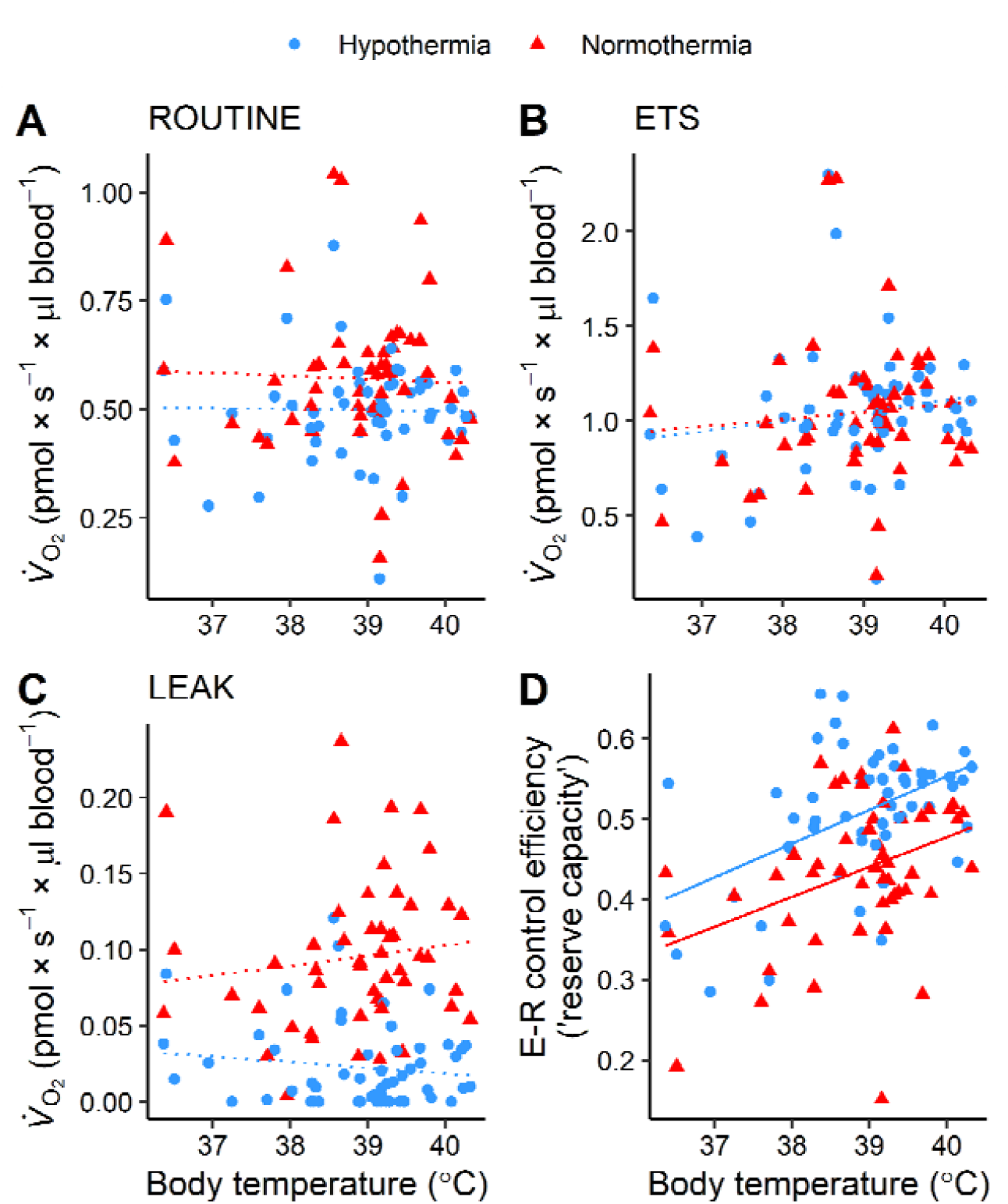
The relationship between mitochondrial respiration traits measured in great tit whole blood and mean body temperature of the individual during the preceding night. Panels A-C show these relationships for ROUTINE, ETS, and LEAK respiration, neither of which were statistically significant as denoted by dotted lines. Panel D) shows the significant relationship with E-R control efficiency, an index of mitochondrial reserve capacity, denoted by the solid lines. Data were collected from wild great tits and respiration was assayed at each of a normothermic (41°C; red triangles) and a hypothermic (35°C; blue circles) thermal state *in vitro*. Sample sizes and statistics are reported in Table S2.

## DISCUSSION

We tested whether plasticity of mitochondrial function could compensate for any reduced cellular respiration rate during rest-phase hypothermia in great tit blood cells. As predicted, endogenous respiration (ROUTINE) was lower in the hypothermic thermal state (Fig. 2A). Surprisingly, this was not a reflection of a general downregulation of cellular respiration during hypothermia, because there was no temperature-dependence of phosphorylating respiration (OXPHOS; in which ATP is produced) (Fig. 2B). Instead, lower ROUTINE was primarily caused by a threefold reduction in LEAK (Fig. 2D). Lack of temperature effects on OXPHOS contrasts previous studies on mammals, where a lowering of assay temperature by 10-12°C to simulate a deep hypothermic state was associated with a 50-70% decrease of both phosphorylating and non-phosphorylating respiration [54-55]. It has been proposed that, in hibernating mammals, active suppression of phosphorylating respiration only becomes noticeable at body temperatures < 30°C [37], which could explain part of the thermal insensitivity of OXPHOS recorded here for a 6°C temperature reduction. If this applies also to birds, a logical conclusion is that passive thermal effects do not influence phosphorylating respiration strongly over the range of temperatures in our study. By contrast, leak respiration showed thermal sensitivity at, or beyond, that recorded in other studies [54-55], but our experimental design does not permit analysis of the contribution of passive thermal and actively regulated effects to this response. Mechanisms aside, the combined actions of thermal insensitivity of OXPHOS and pronounced thermal sensitivity of LEAK meant that the hypothermic thermal state was associated with significant increases in both phosphorylating efficiency (i.e., R-L control efficiency was higher; Fig. 3B) and coupling efficiency of electron transport when the cells were stimulated to work at maximal rate (i.e., E-L coupling efficiency was higher; Fig. 3C), broadly in keeping with results from mammal studies that manipulated assay temperature [54-55]. In fact, nearly all respiration (96%) was directed towards ATP production during hypothermia, and the electron transport system was almost perfectly coupled (0.98%) in this thermal state. It is also interesting to note that we found no effect of simulated hypothermia on maximally stimulated mitochondrial respiration (i.e., ETS; Fig. 2C), suggesting lack of noticeable thermal sensitivity of the electron transport system. As a result, hypothermic cells had higher reserve capacity (Fig. 3A), which could have functional significance in face of a challenge to energy metabolism. Collectively, our study indicates that plasticity of mitochondrial leak respiration combined with thermal insensitivity of phosphorylating and maximal respiration may allow the little bird in winter to secure sufficient ATP production to meet thermoregulatory demands during rest-phase hypothermia. This notion now needs confirmation in other tissues and by *in vivo* studies.

Reduced LEAK and improved phosphorylating and coupling efficiency during hypothermia is in keeping with the hypothesis of biochemical compensation for thermal suppression of cell respiration at low body temperature. However, tighter coupling can potentially result in increased production of reactive oxygen species (ROS) as a by-product of electron transport [21, 56], provided that the resultant increase in protonmotive force is not dissipated through ATP synthase. Unless quenched, ROS cause oxidative stress with resultant macromolecular damage [e.g. 57], which is related to many medical disorders [58]. It is interesting to speculate that a hypothermic bird might have to accept an oxidative cost for the improved oxidative and coupling efficiencies. In this context, our study could provide proximate insights into why oxidative stress has been found to be higher in birds that are more hypothermic at night [59]. However, the role of mitochondria as sources of cellular ROS has been contended [60-62] and empirical evidence suggests that the relationship between body temperature, ATP production and ROS generation is not straightforward [31, 63, 64]. In line with this, even though (pharmacological) uncoupling protects birds from cold-induced oxidative stress [65], the evidence for clear fitness effects thereof is equivocal [66, 67].

We found no strong effects of temperature of the preceding night on mitochondrial respiration traits and flux control ratios, though phosphorylating capacity was the highest after the coldest night-time temperature (Table S1). This contrasts what has been recorded as part of seasonal acclimatisation of blood cell respiration in the same model system [18] and in response to cold exposure in other tissue types [15-17]. Responses in other studies were likely driven by seasonal upregulation of mitochondrial content (e.g. [18]) and so is possible that the period of exposure (one night) and acute nature of the air temperature manipulation was too short or too stochastic to serve as a reliable environmental cue prompting a change to cellular respiratory capacity. It is also possible that seasonal changes in mitochondrial content or respiration rate in the studies above were not causally related to cold temperature. In line with this, cold acclimation did not impact the respiration of liver mitochondria in the lesser hedgehog tenrec (*Echinops telfairi*) [54]. Even so, the difference in short- and long-term responses of mitochondrial respiration coincident with colder environmental temperature suggests it would be interesting to study the temporal resolution of mitochondrial thermal acclimatisation in some detail. For example, it is possible that, in the short term, mitochondrial function responds primarily to a change of tissue temperature in endothermic animals. This could be beneficial because endotherm body temperature is determined by the balance between heat production and heat loss rates and so may be partly or fully independent of air temperature.

On average, there were few effects of night-time body temperature on mitochondrial respiration traits the day after. Similar results were obtained for liver mitochondria in the golden spiny mouse (*Acomys russatus*) measured at a constant assay temperature [68]. Thus, for many of the mitochondrial traits measured here, between-individual variation (which explained 30-70% of total variance in the statistical models; Table S2) did not constrain thermal plasticity. One notable exception was the mitochondrial reserve capacity (i.e., E-R control ratio), which was higher in the hypothermic state (Fig. 3A) but showed a negative relationship with overnight body temperature (Fig. 4). Thus, *in vivo*, more hypothermic birds had mitochondria with *lower* surplus capacity. This invites the speculation that the scope for plastic upregulation of mitochondrial work rate, e.g. in response to sudden environmental change, may be limited by individual differences in the underlying mitochondrial phenotype. It remains to be tested if such constraints on plasticity hinders adaptation to the environment over short- and long temporal scales. Finally, it is interesting to note that the reduction in surplus mitochondrial capacity with decreasing body temperature is well in line with the observation that more hypothermic birds are typically of lower phenotypic quality [e.g., 69-70]. This raises the question of whether rest-phase hypothermia in the little bird in winter is an adaptive response to save energy or if it instead reflects a phenotypic expression enforced by the inability to stay warm.

This study provides new perspectives to our understanding of how cellular and whole animal metabolism interact when the little bird in winter adapts to low temperatures. We conclude that *in vitro* hypothermia was associated with an immediate reduction in proton leak and resultant increased in coupling and oxidative efficiency, which can explain why blood cell mitochondria could maintain phosphorylating respiration despite acute temperature reduction. Nor did low assay temperature suppress maximum working capacity of the electron transport system, leading to increased mitochondrial respiratory scope during hypothermia (i.e., E-R control efficiency was higher, Fig. 3A). However, it is possible that mitochondrial plasticity during rest-phase hypothermia may come at the cost of increased oxidative stress [cf. 59].

Future work should explore this hypothesis by direct measurement of ATP and ROS production, and membrane potential, to conclude on the consequences of changes to the efficiency of phosphorylating respiration for redox balance in normo- and hypothermic birds. Several endotherm studies show that mitochondrial function in blood cells correlates to that in tissues and organs with clearly defined roles in heat production and seasonal acclimatisation (see above), suggesting data presented here may be representative of cellular responses to hypothermia at organismal levels. More broadly, our work aligns with the body of evidence highlighting the utility of blood cell respirometry to gain minimally invasive insight into functional ecology, evolutionary physiology, and gerontology in birds [40, 41, 43,71]. Even so, it is important that future studies formally address the uniformity of mitochondrial plasticity across tissues and organs. Likewise, future investigations should aim to increasingly integrate tissue-based studies with an *in vivo* approach to understand mitochondrial regulation during hypothermia at the organismal level. More broadly, it would be relevant to increase taxonomic and phenotypic coverage of these studies, spanning cold-hardy, non-hypothermic, birds [e.g. 72] to those that routinely use shallow [e.g. 73] or deep torpor [74-76].

## Supporting information

Electronic Supplementary Material 1

## ACKNOWLEDGEMENTS

The authors thank Stefan Nord for managing the Räften feeding stations by which birds for the study were caught. SN also assisted with fieldwork and provided miscellaneous technical support during the study. Two anonymous reviewers provided critical comments that improved the first draft.

## ETHICS

All procedures were approved by the Malmö/Lund Animal Ethics Committee, acting under jurisdiction of the Swedish Board of Agriculture (permit no. 9246-19). Bird ringing was permitted by the Swedish Bird Ringing Centre at Natural History Museum of Sweden (license number 723, to AN).

## DATA AVAILABILITY

Data are deposited in Figshare: 10.6084/m9.figshare.19902277

## REFERENCES

1. Ruf T., Geiser F. 2015 Daily torpor and hibernation in birds and mammals. Biol Rev 90(3), 891–926. (doi:10.1111/brv.12137).

2. Geiser F. 2021 Ecological physiology of daily torpor and hibernation. Cham, Switzerland, Springer Nature Switzerland AG; 317 p.

3. Brodin A. 2007 Theoretical models of adaptive energy management in small wintering birds. Phil Trans R Soc B362(1486), 1857–1871.

4. Nilsson J.F., Nilsson J.-Å., Broggi J., Watson H. 2020 Predictability of food supply modulates nocturnal hypothermia in a small passerine. Biol Lett 16(6), 20200133. (doi:doi:10.1098/rsbl.2020.0133).

5. Nord A., Nilsson J.F., Nilsson J.-Å. 2011 Nocturnal body temperature in wintering blue tits is affected by roost-site temperature and body reserves. Oecologia 167(1), 21–25.

6. Nord A., Nilsson J.F., Sandell M.I., Nilsson J.-Å. 2009 Patterns and dynamics of rest-phase hypothermia in wild and captive blue tits during winter. J Comp Physiol B 179(6), 737–745. (doi:10.1007/s00360-009-0357-1[doi]).

7. Reinertsen R.E., Haftorn S. 1984 The effect of short-time fasting on metabolism and nocturnal hypothermia in the willow tit Parus montanus. J Comp Physiol 154(1), 23–28.

8. Reinertsen R.E., Haftorn S. 1983 Nocturnal hypothermia and metabolism in the willow tit Parus montanus at 63 degrees N. J Comp Physiol 151(2), 109–118.

9. Haftorn S. 1972 Hypothermia of tits in the Arctic winter. Orn Scand 3(2), 153–166.

10. Cooper S.J., Gessaman J.A. 2005 Nocturnal hypothermia in seasonally acclimatized Mountain Chickadees and Juniper Titmice. Condor 107(1), 151–155.

11. Brodin A., Nilsson J.-Å., Nord A. 2017 Adaptive temperature regulation in the little bird in winter: predictions from a stochastic dynamic programming model. Oecologia 185(1), 43–54. (doi:10.1007/s00442-017-3923-3).

12. Welton N.J., Houston A.I., Ekman J., McNamara J.M. 2002 A dynamic model of hypothermia as an adaptive response by small birds to winter conditions. Acta Biotheoretica 50(1), 39–56.

13. Andreasson F., Nord A., Nilsson J.-Å. 2020 Age differences in night-time metabolic rate and body temperature in a small passerine. J Comp Physiol B 190(3), 349–359. (doi:10.1007/s00360-020-01266-5).

14. Nicholls D.G., Ferguson S.J. 2013 Bioenergetics. 4th ed. Cambridge, MA, USA, Academic Press.

15. Liknes E.T., Swanson D.L. 2011 Phenotypic flexibility in passerine birds: Seasonal variation of aerobic enzyme activities in skeletal muscle. J Therm Biol 36(7), 430–436. (doi:http://dx.doi.org/10.1016/j.jtherbio.2011.07.011).

16. Wang Y., Shan S., Zhang H., Dong B., Zheng W., Liu J. 2019 Physiological and biochemical thermoregulatory responses in male Chinese hwameis to seasonal acclimatization: Phenotypic flexibility in a small passerine. Zool Stud 58, e6–e6. (doi:10.6620/ZS.2019.58-06).

17. Zheng W.H., Li M., Liu J.-S., Shao S.-L., Xu X.-J. 2014 Seasonal variation of metabolic thermogenesis in Eurasian tree Sparrows (Passer montanus) over a latitudinal gradient. Physiol Biochem Zool87(5), 704–718. (doi:doi:10.1086/676832).

18. Nord A., Metcalfe N.B., Page J.L., Huxtable A., McCafferty D.J., Dawson N.J. 2021 Avian red blood cell mitochondria produce more heat in winter than in autumn. FASEB J 35(5), e21490. (doi:https://doi.org/10.1096/fj.202100107R).

19. Schaeffer P.J., Villarin J.J., Lindstedt S.L. 2003 Chronic Cold Exposure Increases Skeletal Muscle Oxidative Structure and Function in Monodelphis domestica, a Marsupial Lacking Brown Adipose Tissue. Physiol Biochem Zool76(6), 877–887. (doi:10.1086/378916).

20. Wright K.P., Jr., Hull J.T., Czeisler C.A. 2002 Relationship between alertness, performance, and body temperature in humans. Am J Physiol 283(6), R1370–1377. (doi:10.1152/ajpregu.00205.2002).

21. Divakaruni A.S., Brand M.D. 2011 The Regulation and Physiology of Mitochondrial Proton Leak. Physiology 26(3), 192–205. (doi:10.1152/physiol.00046.2010).

22. Porter R.K., Brand M.D. 1995 Causes of differences in respiration rate of hepatocytes from mammals of different body mass. Am J Physiol 269(5), R1213–1224. (doi:10.1152/ajpregu.1995.269.5.R1213).

23. Rolfe D.F., Brand M.D. 1996 Contribution of mitochondrial proton leak to skeletal muscle respiration and to standard metabolic rate. Am J Physiol 271(4), C1380–1389. (doi:10.1152/ajpcell.1996.271.4.C1380).

24. Stager, M., Cheviron, Z.A. 2020 Is there a role for sarcolipin in avian facultative thermogenesis in extreme cold? Biol Lett 16, 20200078.

25. Hohtola, E. 2004 Shivering thermogenesis in birds and mammals. In Barnes, B.M., Carey, H.V. (eds) Life in the Cold – Mechanisms, adaptation and application, pp. 241–252. Fairbanks: University of Alaska.

26. Teulier, L., Rouanet, J.L., Letexier, D., Romestaing, C., Belouze, M., Rey, B., Duchamp, C., Roussel, D. 2010 Cold-acclimation-induced non-shivering thermogenesis in birds is associated with upregulation of avian UCP but not with innate uncoupling or altered ATP efficiency. J Exp Biol 14, 2476–2482.

27. Nowack, J., Giroud, S., Arnold, W., Ruf, T. 2017 Muscle non-shivering thermogenesis and its role in the evolution of endothermy. Front Physiol 8, 889.

28. Roussel D., Marmillot V., Monternier P.-A., Bourguignon A., Toullec G., Romestaing C., Duchamp C. 2020 Skeletal muscle metabolism in sea-acclimatized king penguins. II. Improved efficiency of mitochondrial bioenergetics. J Exp Biol 223(21), jeb233684. (doi:10.1242/jeb.233684).

29. Bourguignon A., Rameau A., Toullec G., Romestaing C., Roussel D. 2017 Increased mitochondrial energy efficiency in skeletal muscle after long-term fasting: its relevance to animal performance. J Exp Biol 220(13), 2445–2451. (doi:10.1242/jeb.159087).

30. Monternier P.A., Marmillot V., Rouanet J.L., Roussel D. 2014 Mitochondrial phenotypic flexibility enhances energy savings during winter fast in king penguin chicks. J Exp Biol 217(15), 2691–2697. (doi:10.1242/jeb.104505).

31. Boël M., Romestaing C., Duchamp C., Veyrunes F., Renaud S., Roussel D., Voituron Y. 2020 Improved mitochondrial coupling as a response to high mass-specific metabolic rate in extremely small mammals. J Exp Biol 223(5), jeb215558. (doi:10.1242/jeb.215558).

32. Glanville E.J., Seebacher F. 2006 Compensation for environmental change by complementary shifts of thermal sensitivity and thermoregulatory behaviour in an ectotherm. J Exp Biol 209(24), 4869–4877. (doi:10.1242/jeb.02585).

33. Seebacher F. 2005 A review of thermoregulation and physiological performance in reptiles: what is the role of phenotypic flexibility? J Comp Physiol B 175(7), 453–461. (doi:10.1007/s00360-005-0010-6).

34. Bouchard P., Guderley H. 2003 Time course of the response of mitochondria from oxidative muscle during thermal acclimation of rainbow trout, Oncorhynchus mykiss. J Exp Biol 206(Pt 19), 3455–3465. (doi:10.1242/jeb.00578).

35. St-Pierre J., Brand M.D., Boutilier R.G. 2000 The effect of metabolic depression on proton leak rate in mitochondria from hibernating frogs. J Exp Biol 203(9), 1469–1476.

36. Bishop T., St-Pierre J., Brand M.D. 2002 Primary causes of decreased mitochondrial oxygen consumption during metabolic depression in snail cells. Am J Physiol 282(2), R372–R382. (doi:10.1152/ajpregu.00401.2001).

37. Staples J.F., Brown J.C.L. 2008 Mitochondrial metabolism in hibernation and daily torpor: a review. J Comp Physiol B 178(7), 811–827.

38. McKechnie A.E., Lovegrove B.G. 2002 Avian facultative hypothermic responses: A review. Condor 104(4), 705–724.

39. Nord A., Chiriac S., Hasselquist D., Nilsson J.-Å. 2013 Endotoxin injection attenuates rest-phase hypothermia in wintering great tits through the onset of fever. Funct Ecol 27(1), 236–244. (doi:10.1111/1365-2435.12003).

40. Casagrande S., Stier A., Monaghan P., Loveland J.L., Boner W., Lupi S., Trevisi R., Hau M. 2020 Increased glucocorticoid concentrations in early life cause mitochondrial inefficiency and short telomeres. J Exp Biol 223(15), jeb222513. (doi:10.1242/jeb.222513).

41. Stier A., Bize P., Hsu B.-Y., Ruuskanen S. 2019 Plastic but repeatable: rapid adjustments of mitochondrial function and density during reproduction in a wild bird species. Biol Lett 15(11), 20190536. (doi:doi:10.1098/rsbl.2019.0536).

42. Udino E., George J.M., McKenzie M., Pessato A., Crino L., Buchanan K.L., Mariette M.M. 2021 Prenatal acoustic programming of mitochondrial function for high temperatures in an arid-adapted bird. Proc R Soc Lond B 288, 20211893. (doi:doi:10.1098/rspb.2021.1893).

43. Dawson N.J., Salmón P. 2020 Age-related increase in mitochondrial quantity may mitigate a decline in mitochondrial quality in red blood cells from zebra finches (Taeniopygia guttata). Exp Gerontol 133, 110883. (doi:https://doi.org/10.1016/j.exger.2020.110883).

44. Stier A., Romestaing C., Schull Q., Lefol E., Robin J.-P., Roussel D., Bize P. 2017 How to measure mitochondrial function in birds using red blood cells: a case study in the king penguin and perspectives in ecology and evolution. Meth Ecol Evol 8(10), 1172–1182. (doi:10.1111/2041-210x.12724).

45. Stier A., Monaghan P., Metcalfe N.B. 2022 Experimental demonstration of prenatal programming of mitochondrial aerobic metabolism lasting until adulthood. Proc R Soc Lond B 289(1970), 20212679. (doi:doi:10.1098/rspb.2021.2679).

46. Karamercan M.A., Weiss S.L., Villarroel J.P., Guan Y., Werlin E., Figueredo R., Becker L.B., Sims C. 2013 Can peripheral blood mononuclear cells be used as a proxy for mitochondrial dysfunction in vital organs during hemorrhagic shock and resuscitation? Shock 40(6), 476–484. (doi:10.1097/shk.0000000000000026).

47. Petit M., Lewden A., Vézina F. 2014 How does flexibility in body composition relate to seasonal changes in metabolic performance in a small passerine wintering at northern latitude? Physiol Biochem Zool87(4), 539–549. (doi:10.1086/676669).

48. Nord A., Chamkha I., Elmér E. 2022 A whole blood approach improves speed and accuracy when measuring mitochondrial respiration in avian hematocytes. bioRxiv, 2022.2005.2025.493398. (doi:10.1101/2022.05.25.493398).

49. Prinzinger R., Pressmar A., Schleucher E. 1991 Body temperature in birds. Comp Biochem Physiol A 99(4), 499–506.

50. Gnaiger E. 2020 Mitochondrial pathways and respiratory control: An Introduction to OXPHOS Analysis. 5th ed.

51. Bates D., Maechler M., Bolker B., Walker S. 2015 Fitting linear mixed-effects models using lme4. J Stat Soft 67, 1–48.

52. Halekoh, U., Højsgaard, S. 2014 A Kenward-Roger approximation and parametric bootstrap methods for tests in linear mixed models – The R package pbkrtest. J Stat Soft 59(9), 1–32.

53. Lenth R.V. 2019 emmeans: Estimated marginal means, aka least-squares means. R package version 1.3.3. https://CRAN.R-project.org/package=emmeans.

54. Polymeropoulos, E.T., Oelkrug, R., Jastroch, M. 2017 Mitochondrial proton leak compensates for reduced oxidative power during frequent hypothermic events in a protoendothermic mammal, Echinops telfairi. Front Physiol 8, 909.

55. Dufour, S., Rousse, N., Canioni, P, Diolez, P. 1996 Top-down control analysis of temperature effect on oxidative phosphorylation. Biochem J 314, 743–751.

56. Demine S., Renard P., Arnould T. 2019 Mitochondrial uncoupling: A key controller of biological processes in physiology and diseases. Cells 8(8), 795. (doi:10.3390/cells8080795).

57. Brand M.D., Buckingham J.A., Esteves T.C., Green K., Lambert A.J., Miwa S., Murphy M.P., Pakay J.L., Talbot D.A., Echtay K.S. 2004 Mitochondrial superoxide and aging: uncoupling-protein activity and superoxide production. Biochem Soc Symp (71), 203–213. (doi:10.1042/bss0710203).

58. Grasemann H., Holguin F. 2021 Oxidative stress and obesity-related asthma. Paed Resp Rev 37, 18–21. (doi:10.1016/j.prrv.2020.05.004).

59. Zagkle E., Grosiak M., Bauchinger U., Sadowska E.T. 2020 Rest-phase hypothermia reveals a link between aging and oxidative stress: A novel hypothesis. Front Physiol 11, 575060. (doi:10.3389/fphys.2020.575060).

60. Zhang Y., Wong H.S. 2021 Are mitochondria the main contributor of reactive oxygen species in cells? J Exp Biol 224(5), eb221606. (doi:10.1242/jeb.221606).

61. Mookerjee S.A., Divakaruni A.S., Jastroch M., Brand M.D. 2010 Mitochondrial uncoupling and lifespan. Mech Ageing Devel 131(7-8), 463–472. (doi:10.1016/j.mad.2010.03.010).

62. Brown G.C., Borutaite V. 2012 There is no evidence that mitochondria are the main source of reactive oxygen species in mammalian cells. Mitochondrion 12(1), 1–4. (doi:https://doi.org/10.1016/j.mito.2011.02.001).

63. Roussel D., Voituron Y. 2020 Mitochondrial costs of being hot: Effects of acute thermal change on liver bioenergetics in toads (Bufo bufo). Front Physiol 11, 153. (doi:10.3389/fphys.2020.00153).

64. Boël M., Veyrunes F., Durieux A.-C., Freyssenet D., Voituron Y., Roussel D. 2022 Does high mitochondrial efficiency carry an oxidative cost? The case of the African pygmy mouse (Mus mattheyi). Comp Biochem Physiol A 264, 111111. (doi:https://doi.org/10.1016/j.cbpa.2021.111111).

65. Stier A., Massemin S., Criscuolo F. 2014 Chronic mitochondrial uncoupling treatment prevents acute cold-induced oxidative stress in birds. J Comp Physiol B 184(8), 1021–1029. (doi:10.1007/s00360-014-0856-6).

66. Stier A., Bize P., Roussel D., Schull Q., Massemin S., Criscuolo F. 2014 Mitochondrial uncoupling as a regulator of life-history trajectories in birds: an experimental study in the zebra finch. J Exp Biol 217(19), 3579–3589. (doi:10.1242/jeb.103945).

67. Stier A., Bize P., Massemin S., Criscuolo F. 2021 Long-term intake of the illegal diet pill DNP reduces lifespan in a captive bird model. Comp Biochem Physiol C 242, 108944. (doi:https://doi.org/10.1016/j.cbpc.2020.108944).

68. Grimpo, K., Kutschke, M., Kastl, A., Meyer, C.W., Heldmaier, G., Exner, C., Jastroch, M. 2014 Metabolic depression during warm torpor in the Golden spiny mouse (Acomys russatus) does not affect mitochondrial respiration and hydrogen peroxide release. Comp Biochem Physiol A 167, 7–14.

69. Nord, A., Nilsson, J.F., Sandell, M.I., Nilsson, J.-Å. 2009 Patterns and dynamics of rest-phase hypothermia in wild and captive blue tits during winter. J Comp Physiol B 179(6), 737–745.

70. Nord, A., Nilsson, J.F., Nilsson, J.-Å. 2011 Nocturnal body temperature in wintering blue tits is affected by roost-site temperature and body reserves. Oecologia 167(1), 21–25.

71. Brodin, A., Nilsson, J.-Å., Nord, A. 2017 Adaptive temperature regulation in the little bird in winter: predictions from a stochastic dynamic programming model. Oecologia 185(1), 43–54.

72. Stier A., Bize P., Schull Q., Zoll J., Singh F., Geny B., Gros F., Royer C., Massemin S., Criscuolo F. 2013 Avian erythrocytes have functional mitochondria, opening novel perspectives for birds as animal models in the study of ageing. Frontiers in Zoology 10(1), 33. (doi:10.1186/1742-9994-10-33).

73. Nord A., Folkow L.P. 2018 Seasonal variation in the thermal responses to changing environmental temperature in the world’s northernmost land bird. J Exp Biol 221(1), jeb171124. (doi:10.1242/jeb.171124).

74. Romano A.B., Hunt A., Welbergen J.A., Turbill C. 2019 Nocturnal torpor by superb fairy-wrens: a key mechanism for reducing winter daily energy expenditure. Biol Lett 15(6), 20190211. (doi:doi:10.1098/rsbl.2019.0211).

75. Hiebert S.M. 1990 Energy costs and temporal organization of torpor in the rufous hummingbird (Selasphorus rufus). Physiol Zool 63, 1082–1097.

76. Brigham R.M. 1992 Daily torpor in a free-ranging goatsucker, the common poorwill (Phalaenoptilus nuttallii). Physiol Zool 65(2), 457–472. (doi:10.2307/30158263).

