## Supplementary material for "Plasticity of mitochondrial function safeguards phosphorylating respiration during *in vitro* simulation of rest-phase hypothermia": Electronic Supplementary Material 1

**Electronic Supplementary Material for**

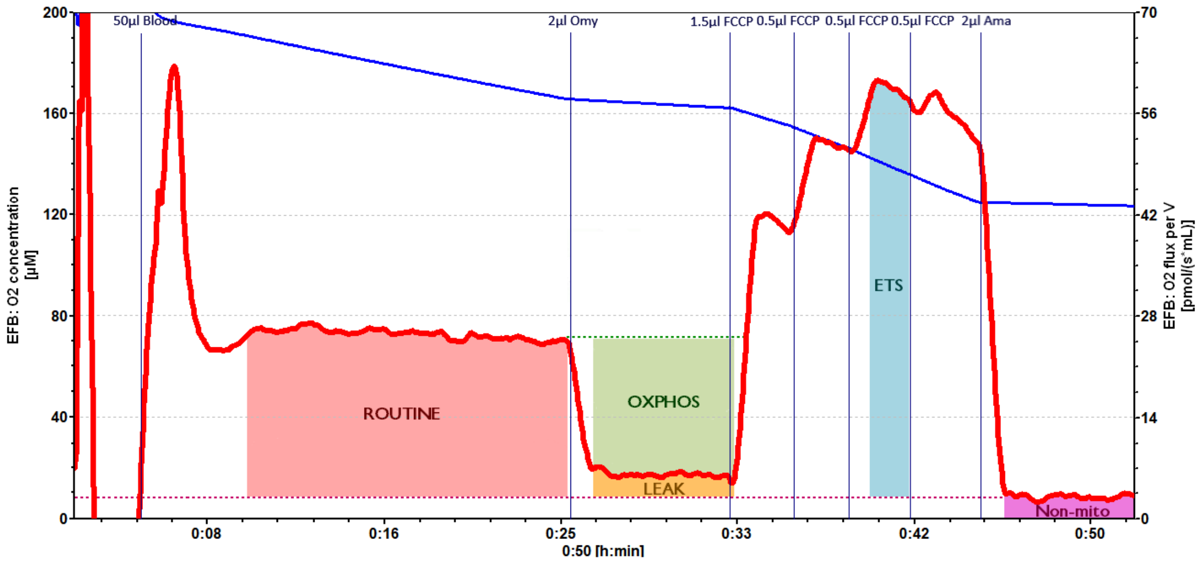


**Figure S1**. Representative experiment testing mitochondrial respiration in whole blood of a winter-adapted great tit (*Parus major*, individual 2KV07141) when measured at a normothermic daytime bird body temperature (41 °C). The blue line shows oxygen ($O_2$) concentration in the test chamber, and the red line shows $O_2$ consumption by the mitochondria on endogenous substrates (ROUTINE), after the addition of ATP synthase inhibitor oligomycin (Omy) (OXPHOS), when maximally stimulated by the mitochondrial uncoupler FCCP (carbonyl cyanide-p-trifluoro-methoxyphenyl-hydrazone) (ETS), and after inhibition by the cytochrome c reductase inhibitor antimycin A (Ama). LEAK refers to the part of respiration devoted to compensating for protons escaping over the inner mitochondrial membrane. "Non-mito" refers to non-mitochondrial respiration, i.e., oxygen consumption by the sample (such as binding of $O_2$ by haemoglobin) that is not due to mitochondrial use. Respiration states are defined in Table 1 in the main manuscript and were superimposed onto the raw data for clarity.

**Table S1.** Parameter estimates, sample sizes, test statistics, degrees of freedom, and resultant *P*-values from linear models testing how a sudden change in night temperature affected mitochondrial respiration parameters and respiratory control ratios in normothermic great tits the morning after. Model-estimated means are presented ± s.e.m. Significant effects (i.e., $P \leq 0.05$) are presented in bold font, and effects for which $0.05 > P < 0.10$ are given in italics.

| Model and Variable | Estimate ± s.e.m. | *n* | d.f. | *F* | *P* |
|---|---|---|---|---|---|
| **ROUTINE (pmol $O_2$ $s^{-1}$ $\mu l^{-1}$)** | | 52 | | | |
| Night temp (-15°C, 5°C or 25°C) | | | 2, 49 | 0.51 | 0.607 |
| Night temp = -15°C | 0.565±0.041 | 17 | | | |
| Night temp = 5°C | 0.546±0.058 | 17 | | | |
| Night temp = 25°C | 0.602±0.057 | 18 | | | |
| **OXPHOS (pmol $O_2$ $s^{-1}$ $\mu l^{-1}$)** | | 52 | | | |
| Night temp (-15°C, 5°C or 25°C) | | | 2, 49 | 0.79 | 0.461 |
| Night temp = -15°C | 0.487±0.034 | | | | |
| Night temp = 5°C | 0.442±0.047 | | | | |
| Night temp = 25°C | 0.498±0.047 | | | | |
| **ETS (pmol $O_2$ $s^{-1}$ $\mu l^{-1}$)** | | 52 | | | |
| Night temp (-15°C, 5°C or 25°C) | | | 2, 49 | 0.99 | 0.376 |
| Night temp = -15°C | 0.981±0.090 | 17 | | | |
| Night temp = 5°C | 0.993±0.127 | 17 | | | |
| Night temp = 25°C | 0.134±0.125 | 18 | | | |
| **LEAK (pmol $O_2$ $s^{-1}$ $\mu l^{-1}$)** | | 52 | | | |
| Night temp (-15°C, 5°C or 25°C) | | | 2, 49 | 1.56 | 0.220 |
| Night temp = -15°C | 0.078±0.012 | 17 | | | |
| Night temp = 5°C | 0.103±0.016 | 17 | | | |
| Night temp = 25°C | 0.104±0.016 | 18 | | | |
| **E-R control efficiency (‘mitochondrial reserve capacity’)** | | 52 | | | |
| Night temp (-15°C, 5°C or 25°C) | | | 2, 49 | 1.60 | 0.213 |
| Night temp = -15°C | 0.408±0.023 | 17 | | | |
| Night temp = 5°C | 0.419±0.033 | 17 | | | |
| Night temp = 25°C | 0.462±0.032 | 18 | | | |
| **R-L control efficiency (‘phosphorylating capacity’)** | | 52 | | | |
| Night temp (-15°C, 5°C or 25°C) | | | 2, 49 | 2.92 | *0.063* |
| Night temp = -15°C | 0.858±0.014 | 17 | | | |
| Night temp = 5°C | 0.811±0.020 | 17 | | | |
| Night temp = 25°C | 0.834±0.019 | 18 | | | |
| **E-L coupling efficiency (‘tightness of electron transport’)** | | 52 | | | |
| Night temp (-15°C, 5°C or 25°C) | | | 2, 49 | 1.90 | 0.160 |
| Night temp = -15°C | 0.915±0.009 | 17 | | | |
| Night temp = 5°C | 0.892±0.013 | 17 | | | |
| Night temp = 25°C | 0.911±0.012 | 18 | | | |

**Table S2.** Parameter estimates, sample sizes, degrees of freedom, test statistics and resultant *P*-values from linear mixed effects models used to investigate how mitochondrial respiration parameters and respiratory control ratios in great tit whole blood was affected by a change from a normothermic (41 °C) to a hypothermic (35 °C) thermal state. Model-estimated means are presented ± s.e.m. Significant effects (i.e., $P \leq 0.05$) are presented in bold font, and effects for which $0.05 > P < 0.10$ are given in italics. Degrees of freedom (d.f.) were calculated using the Kenward-Roger approximation. The test statistic is *F* for all fixed effects and the likelihood ratio (LRT) for the random term. Note that *P*-values for main effects were not provided when the interaction was significant. *Main effect to be interpreted with caution due to involvement in an interaction with mean nightly body temperature.

| Model and Variable | Estimate ± s.e.m. | $n_{hyp}$ / $n_{norm}$ | d.f. | *F / LRT* | *P* | $\sigma^2_{Bird}$ \| $\sigma^2_{Residual}$ |
|---|---|---|---|---|---|---|
| **ROUTINE (pmol $O_2$ $s^{-1}$ $\mu l^{-1}$)** | | 57 / 52 | | | | |
| Body temperature × Thermal state | | | 1, 53.0 | 0.04 | 0.850 | |
| Body temperature (°C) | -0.002±0.019 | | 1, 54.5 | 0.02 | 0.903 | |
| Thermal state | | | 1, 53.1 | 22.16 | **< 0.0001** | |
| Hypothermia (35°C) | 0.499±0.019 | | | | | |
| Normothermia (41°C) | 0.567±0.019 | | | | | |
| Bird ID (random) | | | 1 | 41.23 | **< 0.0001** | 0.0154 \| 0.0055 |
| **OXPHOS (pmol $O_2$ $s^{-1}$ $\mu l^{-1}$)** | | 57 / 52 | | | | |
| Body temperature × Thermal state | | | 1, 53.3 | 0.29 | 0.592 | |
| Body temperature (°C) | -0.002±0.016 | | 1, 54.5 | 0.02 | 0.879 | |
| Thermal state | | | 1, 53.3 | 0.11 | 0.744 | |
| Hypothermia (35°C) | 0.476±0.017 | | | | | |
| Normothermia (41°C) | 0.472±0.017 | | | | | |
| Bird ID (random) | | | 1 | 34.74 | **< 0.0001** | 0.0109 \| 0.0046 |
| **ETS (pmol $O_2$ $s^{-1}$ $\mu l^{-1}$)** | | 57 / 52 | | | | |
| Body temperature × Thermal state | | | 1, 51.7 | 0.10 | 0.748 | |
| Body temperature (°C) | 0.054±0.048 | | 1, 54.2 | 1.25 | 0.268 | |
| Thermal state | 1.050±0.047 | | 1, 52.4 | 0.33 | 0.570 | |
| Hypothermia (35°C) | 1.040±0.048 | | | | | |
| Normothermia (41°C) | | | | | | |
| Bird ID (random) | | | 1 | 93.27 | **< 0.0001** | 0.1133 \| 0.0109 |
| **LEAK (pmol $O_2$ $s^{-1}$ $\mu l^{-1}$)** | | 57 / 52 | | | | |
| Body temperature × Thermal state | | | 1, 54.7 | 3.16 | *0.081* | |
| Body temperature (°C) | 0.000±0.005 | | 1, 54.3 | 0.00 | 0.971 | |
| Thermal state | | | 1, 54.2 | 173.24 | **< 0.0001** | |
| Hypothermia (35°C) | 0.023±0.005 | | | | | |
| Normothermia (41°C) | 0.096±0.005 | | | | | |
| Bird ID (random) | | | 1 | 14.28 | **0.0002** | 0.0007 \| 0.0008 |
| **E-R control efficiency (‘mitochondrial reserve capacity’)** | | 57 / 52 | | | | |
| Body temperature × Thermal state | | | 1, 54.1 | 0.02 | 0.882 | |
| Body temperature (°C) | 0.041±0.010 | | 1, 54.5 | 16.62 | **0.0001** | |
| Thermal state | | | 1, 53.7 | 51.34 | **< 0.0001** | |
| Hypothermia (35°C) | 0.505±0.011 | | | | | |
| Normothermia (41°C) | 0.434±0.011 | | | | | |
| Bird ID (random) | | | 1 | 21.79 | **< 0.0001** | 0.0038 \| 0.0026 |
| **R-L control efficiency (‘phosphorylating capacity’)** | | 57 / 52 | | | | |
| Body temperature × Thermal state | | | 1, 55.5 | 4.57 | **0.037** | |
| Hypothermia (35°C) | 0.006±0.007 | | | | | |
| Normothermia (41°C) | -0.012±0.008 | | | | | |
| Body temperature (°C) | | | | | | |
| Thermal state | | | | | | |
| Hypothermia (35°C) | 0.956±0.007* | | | | | |
| Normothermia (41°C) | 0.834±0.007* | | | | | |
| Bird ID (random) | | | 1 | 7.44 | **0.006** | 0.0011 \| 0.0018 |

| | | | | | | |
|---|---|---|---|---|---|---|
| **E-L coupling efficiency (‘tightness of electron transport’)** | | 57 / 52 | | | | |
| Body temperature × Thermal state | | | 1, 56.0 | 0.83 | 0.365 | |
| Body temperature (°C) | 0.003±0.004 | | 1, 53.0 | 0.72 | 0.400 | |
| Thermal state | | | 1, 55.0 | 194.57 | **< 0.0001** | |
| Hypothermia (35°C) | 0.978±0.004 | | | | | |
| Normothermia (41°C) | 0.907±0.004 | | | | | |
| Bird ID (random) | | | 1 | 4.02 | **0.045** | 0.0003 \| 0.0007 |
